# Numericware i: Identical in state matrix calculator

**DOI:** 10.1101/075267

**Authors:** Bongsong Kim, William Beavis

## Abstract

Herein we introduce software, Numericware i to compute a matrix consisting of all pairwise identical in state (IIS) coefficients from genotypic data. Since the emergence of high throughput technology for genotyping, calculating an IIS matrix between many pairs of entities has required large computer memory and lengthy processing times. Numericware i addresses these limitations with two algorithmic methods: multithreading and forward chopping. The multithreading feature allows computational routines to concurrently run on multiple CPU processors. The forward chopping addresses memory limitations by dividing the genotypic data into appropriately sized subsets. Numericware i allows researchers who need to estimate an IIS matrix for big genotypes to use typical laptop/desktop computers. For comparison with different software, we calculated genetic relationship matrices using Numericware i, SPAGeDi and TASSEL with the same small-sized data set. Numericware i measured kinship coefficients between zero and two, while the matrices from SPAGeDi and TASSEL produced different ranges of values, including negative values. The Pearson correlation coefficient between the matrices from Numericware i and TASSEL was high at 0.993, while SPAGeDi rarely showed correlation with Numericware i (0.088) and TASSEL (0.087). To compare the capacity with high dimensional data, we applied the three software to a simulated data set consisted of 500 entities by 1,000,000 SNPs. Numericware i spent 71 minutes using seven CPU cores on a laptop (DELL LATITUDE E6540), while SPAGeDi and TASSEL failed to start. Numericware i is freely available for Windows and Linux under CC-BY license at https://figshare.com/s/f100f33a8857131eb2db.

## Introduction

The inbreeding coefficient, kinship coefficient, and identical by descent (IBC) coefficient are central parameters in population genetics (Cockerham and Weir, 1983). By definition, the inbreeding coefficient refers to a proportion that a pair of alleles in an entity are identical by descent (Wright, 1922), and the kinship coefficient between two entities is equal to twice of the inbreeding coefficient for their virtual offspring (Emik and Terrill, 1949). The kinship coefficient is a conventional indicator to represent genetic relatedness among entities in a population, for which pedigree records are available. Emik and Terrill (1949) provided a systematic method for calculating a numerator relationship matrix (NRM) that displays kinship coefficients among every pair of entities in a population. Because their method uses pedigree records, the results are used to infer genetic relatedness from the genealogical perspective.

High throughput technologies for genomic assays provide abundant DNA profile. This can replace the pedigree records for measuring genetic relatedness. Some references (Bernardo, 2002) introduced a method for computing an identical in state matrix (IIS matrix). The IIS matrix ignores the genetic origin of the alleles, and is thoroughly based on allelic states throughout the genomes. For some applications, it may be more useful than the NRM. Although the concept about the IIS matrix is general, surprisingly the method is not widely used. The method for computing the IIS matrix is simple, but computing workloads are notoriously heavy. Therefore, obtaining an IIS matrix may require extensive data management and parallel computing techniques. In this paper, we present software, referred to as Numericware i, which will compute the IIS matrix using any set of processors available on typical laptop/desktop computers.

## Identical in state coefficient

Given codominant marker data, the IIS coefficients can be obtained based on marker genotypes. The formula for computing the IIS coefficient is as follows:

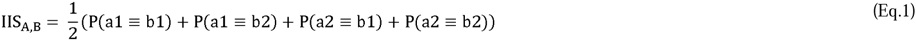

where IIS_A,B_, = the identical in state coefficient between A and B; a1, a2 = a pair of alleles for A; b1, b2 = a pair of alleles for B; P(a1≡b1) = the probability that a1 and b1 are homozygous; P(a1≡b2) = the probability that a1 and b2 are homozygous; P(a2≡b1) = the probability that a2 and b1 are homozygous; P(a2≡b2) = the probability that a2 and b2 are homozygous.

As a counterpart of the kinship coefficient, the IIS coefficient ranges between zero and two. Like the kinship coefficient = twice the inbreeding coefficient, the IIS coefficient = twice the homozygote coefficient (HC). Thus, the HC can be calculated as follows (Bernardo, 2002):

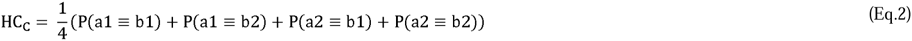

where C = the progeny of A and B; HC_C_ = the homozygous coefficient for C; a1, a2 = a pair of alleles for A; b1, b2 = a pair of alleles for B; P(a1≡b1) = the probability that a1 and b1 are homozygous; P(a1≡b2) = the probability that a1 and b2 are homozygous; P(a2≡b1) = the probability that a2 and b1 are homozygous; P(a2≡b2) = the probability that a2 and b2 are homozygous.

Depending on the central limit theorem, greater numbers of markers provide better representation for the HC and IIS coefficients. As high throughput methods for obtaining DNA fingerprints are becoming cheaper, the dimensions of genotypic data sets are rapidly growing. The amount of computing workload can be represented as:

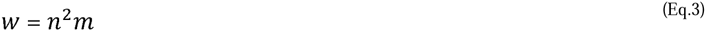

where *w* = the amount of computational workload; *n* = the number of entities in a population; *m* = the number of markers

According to Eq. 3, the growing dimension of genotypic data generates two computational challenges: (1) long computational times; (2) shortage of computer memory. This situation makes the computation of the IIS matrix challenging.

## Functionality of Numericware i

Numericware i is written in C++. The software has a simple user interface, provides both high functionality and ease of use, and enables typical laptop/desktop computers to handle large genotypic data sets using all available system resources. The software provides two special functionalities: multithreading and forward chopping. The multithreading function enables the computer to distribute the workload into multiple CPU cores. Forward chopping is a method that chops a large size of genotypic data into multiple pieces that will not overextend memory allocations. The algorithm of the forward chopping method is as follows:

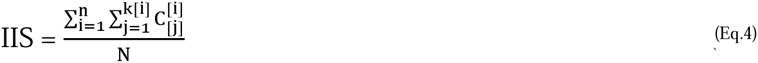

where IIS = the IIS coefficient; n = the number of chopped subsets; i = the loop variable referring to a chopped subset number; j = the loop variable referring to a locus within a chopped subset; k[i] = the number of loci in the i'th chopped subset; 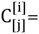 the IIS coefficient in the j'th locus within the i'th subset; N = the total number of loci.

### Numericware i provides users with further conveniences

1. The IIS matrix computation for a haplotype: this procedure allows to compute the IIS matrix with a subset of genomic data.
2. Genotypic data integrity checking: this procedure helps prevent a failure in the middle of analysis by checking the integrity of data at the beginning of work.
3. Genotypic score summary: this provides users with an overview on genotypic scores.
4. Supporting multiple data formats: this significantly reduces an extra work for formatting the input data. Numericware i accepts the following formats: alphanumeric, a pair of SNPs, and IUPAC.
5. Inverting the IIS matrix: the inverted matrix can be directly applied to the best linear unbiased prediction (BLUP) procedure. As an inverting method, the Cholesky decomposition algorithm is implemented.
6. Transposing the genotypic data.

If missing genotypic scores are found, Numericware i skips the missing loci, assuming that the number of non-missing allelic scores is sufficient to represent the genome in a large data set. Therefore, an imputation of missing data points is not required.

## Application of the IIS matrix

HC index: a diagonal value for an entity, A, (IIS_A,A_) in an IIS matrix implies the HC_A_ as IIS_A,A_ = 1 + HC_A_, and the IIS coefficient between two entities represents twice the HC for their virtual offspring. These properties can help control homozygote level of new entities that will be produced in a breeding program.

BLUP: the IIS matrix is superior to the NRM in practicing the BLUP for the following reasons. First, the IIS matrix is complete with coefficients for all pairs of genotypes. In contrast, the NRM includes an identity matrix for a base population. The identity matrix will result in an underestimation of kinship coefficients within the NRM. Second, the IIS matrix provides an objective measure about genetic relatedness that is limited only by the limitations of the marker technology, while the NRM utilizes statistical expectations for IBD. In order for BLUP to provide unbiased solutions caused by genetic relatedness, the IIS matrix can be used as an alternative to the NRM. Since genome wide association mapping (GWAS) and genomic prediction (GP) utilize BLUP methods, the IIS matrix can be used for these applications. The IIS matrix may be more useful for plant breeding than animal breeding since the pedigree records for plants are often unknown and imprecise (Bauer et al, 2006). As Numericware i supports computing haplotype IIS matrix, incorporating a haplotype IIS matrix for a genomic region of interest into the BLUP is possible. Previous studies reported that the BLUP practices using genome-based relationship matrices outperformed those using the NRM (Colleau, 2002; Habier et al, 2007; VanRaden, 2008; Legarra et al, 2009; Endelman and Jannink, 2012; Müller et al, 2015). The IIS matrix will be worth using for the BLUP.

## Negative effect of marker screen to IIS matrix

Genotypic marker technologies often produce DNA markers that have been screened for frequencies of polymorphic alleles. Therefore, many markers might have been filtered out. Marker screening will create an ascertainment bias in the IIS matrix because the filtered non-polymorphic markers must also be informative in representing the genomic state among entities. Therefore, it is recommended that all markers be used for estimating the IIS matrix.

## Similarity between IBD and IIS

Between the IIS matrix and NRM, the range of values is the same between zero and two, but the basic concepts are slightly different. If it is assured that any identical alleles from different entities were generated not independently but inherently, the IIS and IBD coefficients should be equal. We hypothetically assume that mutations mainly create genetic diversity, and that common mutations in a population mostly happened a single time, and were shared inherently since the probability that the same mutations coincidentally happened on the same spots of multiple entities might be extremely low. If this assumption is true, the IIS matrix can represent the true kinship matrix.

## Comparison of results from Numericware i, SPAGeDi, and TASSEL

Henderson (1975) suggested to plug the NRM into the BLUP for correcting a bias caused by genealogical relationship among entities in a population. Assuming that the kinship and IIS coefficients are very close, the precision of NRM can be substantially increased by replacing the identity matrix for the base population by the IIS matrix. Meanwhile, popular software, SPAGeDi (Hardy and Vekemans, 2002) and TASSEL (Bradbury et al, 2007), implement different algorithms for calculating the genetic relationship matrix, causing their results to have different characteristics such as negative elemental values (SPAGeDi and TASSEL), or mono-diagonal values (SPAGeDi with zero). Thus, the resulting matrix from SPAGeDi and TASSEL cannot replace the identity matrix within the NRM. For comparison, we applied Numericware i, SPAGeDi, and TASSEL to the same data set in small size. Their output matrices are shown in Supplementary tables 1 to 3. Pearson correlation coefficients among the three tables (Table) indicate that the results from Numericware i and TASSEL are highly correlated at 0.992, while the result from SPAGeDi rarely shows correlation with the results from Numericware i (0.088) and TASSEL (0.087). This illustrates that the genetic relationship matrices from Numericware i and TASSEL are substantially comparable, while SPAGeDi is not.

**Table.** Pearson correlation coefficients among results from Numericware i, SPAGeDi, and TASSEL with the same genotypic data.

|  | Numericware i | SPAGeDi | TASSEL |
| --- | --- | --- | --- |
| Numericware i | 1 | 0.088 | 0.992 |
| SPAGeDi | 0.088 | 1 | 0.087 |
| TASSEL | 0.992 | 0.087 | 1 |

## Performance

In our test, Numericware i completed computing an IIS matrix with a simulated data set sized as 500 entities by 1,000,000 SNPs for 71 minutes using seven CPU cores on a laptop (DELL LATITUDE E6540). For this test, the whole data was chopped into three pieces to circumvent the memory limit of 8 GB, while SPAGeDi and TASSEL failed with the same data due to the limitation of memory.

## Conclusion

The IIS matrix can be useful for versatile purposes, e.g. measuring the HC for an entity to itself; predicting the HCs for offspring; measuring the genetic relatedness among entities in a population; and practicing the BLUP. Thus, Numericware i can be an essential tool for breeding programs. The multithreading and forward chopping methods remarkably reduce the computing burden, reduce computing time, and allow the computation of extremely big data even with typical laptops or desktop computers. In contrast, analyses with other software are often limited by the physical memory size, and only a single CPU is supported. Furthermore, an easy-to-use interface minimizes users to refer to the user manual, and allows them to quickly get familiar with the software.

## Availability

Numericware i is freely available for Windows and Linux under CC-BY license, and can be downloaded from https://figshare.com/s/f100f33a8857131eb2db.

---

**Supplementary table 1.** Identical in state matrix computed using Numericware i. All elemental values range between zero and two.

|  | ID1 | ID2 | ID3 | ID4 | ID5 | ID6 | ID7 | ID8 | ID9 | ID10 | ID11 | ID12 | ID13 | ID14 | ID15 | ID16 | ID17 | ID18 | ID19 | ID20 |
| --- | --- | --- | --- | --- | --- | --- | --- | --- | --- | --- | --- | --- | --- | --- | --- | --- | --- | --- | --- | --- |
| ID1 | 1.24473 | 0.50338 | 0.52435 | 0.49156 | 0.48333 | 0.49666 | 0.49272 | 0.51069 | 0.48373 | 0.51577 | 0.50724 | 0.48101 | 0.49443 | 0.4927 | 0.46581 | 0.48597 | 0.47917 | 0.4836 | 0.48667 | 0.49606 |
| ID2 | 0.50338 | 1.23184 | 0.49886 | 0.49829 | 0.4786 | 0.50282 | 0.51746 | 0.51471 | 0.49886 | 0.51767 | 0.52599 | 0.50962 | 0.50564 | 0.52386 | 0.49378 | 0.52043 | 0.5234 | 0.50345 | 0.49719 | 0.50795 |
| ID3 | 0.52435 | 0.49886 | 1.22556 | 0.5 | 0.51251 | 0.51524 | 0.49205 | 0.51313 | 0.47143 | 0.49083 | 0.51873 | 0.49659 | 0.51135 | 0.504 | 0.50456 | 0.49543 | 0.48443 | 0.51095 | 0.50962 | 0.48744 |
| ID4 | 0.49156 | 0.49829 | 0.5 | 1.22222 | 0.47968 | 0.5079 | 0.48473 | 0.51873 | 0.48687 | 0.52229 | 0.48253 | 0.50903 | 0.4887 | 0.48636 | 0.50057 | 0.49489 | 0.54176 | 0.50859 | 0.50899 | 0.49943 |
| ID5 | 0.48333 | 0.4786 | 0.51251 | 0.47968 | 1.24182 | 0.49666 | 0.51175 | 0.48822 | 0.49888 | 0.50396 | 0.50501 | 0.4972 | 0.50558 | 0.52025 | 0.52741 | 0.52352 | 0.50227 | 0.49319 | 0.49278 | 0.51179 |
| ID6 | 0.49666 | 0.50282 | 0.51524 | 0.5079 | 0.49666 | 1.26399 | 0.50056 | 0.49551 | 0.50843 | 0.50113 | 0.50834 | 0.4933 | 0.48608 | 0.53371 | 0.48599 | 0.48086 | 0.48702 | 0.51359 | 0.5061 | 0.50901 |
| ID7 | 0.49272 | 0.51746 | 0.49205 | 0.48473 | 0.51175 | 0.50056 | 1.23729 | 0.49607 | 0.49324 | 0.50566 | 0.47263 | 0.49048 | 0.48498 | 0.48142 | 0.53471 | 0.50169 | 0.49887 | 0.48804 | 0.5167 | 0.49606 |
| ID8 | 0.51069 | 0.51471 | 0.51313 | 0.51873 | 0.48822 | 0.49551 | 0.49607 | 1.23617 | 0.4949 | 0.46298 | 0.48597 | 0.52531 | 0.49606 | 0.47571 | 0.50676 | 0.51925 | 0.4886 | 0.4943 | 0.55387 | 0.49831 |
| ID9 | 0.48373 | 0.49886 | 0.47143 | 0.48687 | 0.49888 | 0.50843 | 0.49324 | 0.4949 | 1.24255 | 0.49205 | 0.48992 | 0.49268 | 0.51798 | 0.50849 | 0.49718 | 0.47401 | 0.50741 | 0.49715 | 0.49777 | 0.49095 |
| ID10 | 0.51577 | 0.51767 | 0.49083 | 0.52229 | 0.50396 | 0.50113 | 0.50566 | 0.46298 | 0.49205 | 1.26389 | 0.50675 | 0.52316 | 0.52198 | 0.49036 | 0.52216 | 0.50796 | 0.50517 | 0.50919 | 0.4955 | 0.49886 |
| ID11 | 0.50724 | 0.52599 | 0.51873 | 0.48253 | 0.50501 | 0.50834 | 0.47263 | 0.48597 | 0.48992 | 0.50675 | 1.24684 | 0.5067 | 0.52892 | 0.49381 | 0.5106 | 0.47919 | 0.46546 | 0.45598 | 0.48556 | 0.52301 |
| ID12 | 0.48101 | 0.50962 | 0.49659 | 0.50903 | 0.4972 | 0.4933 | 0.49048 | 0.52531 | 0.49268 | 0.52316 | 0.5067 | 1.24974 | 0.47826 | 0.49944 | 0.48989 | 0.51748 | 0.47839 | 0.50342 | 0.50723 | 0.49211 |
| ID13 | 0.49443 | 0.50564 | 0.51135 | 0.4887 | 0.50558 | 0.48608 | 0.48498 | 0.49606 | 0.51798 | 0.52198 | 0.52892 | 0.47826 | 1.26399 | 0.52918 | 0.51175 | 0.49157 | 0.49548 | 0.49489 | 0.52722 | 0.52301 |
| ID14 | 0.4927 | 0.52386 | 0.504 | 0.48636 | 0.52025 | 0.53371 | 0.48142 | 0.47571 | 0.50849 | 0.49036 | 0.49381 | 0.49944 | 0.52918 | 1.27021 | 0.50788 | 0.50113 | 0.50171 | 0.47722 | 0.49665 | 0.49548 |
| ID15 | 0.46581 | 0.49378 | 0.50456 | 0.50057 | 0.52741 | 0.48599 | 0.53471 | 0.50676 | 0.49718 | 0.52216 | 0.5106 | 0.48989 | 0.51175 | 0.50788 | 1.24047 | 0.51858 | 0.49321 | 0.50227 | 0.49499 | 0.48647 |
| ID16 | 0.48597 | 0.52043 | 0.49543 | 0.49489 | 0.52352 | 0.48086 | 0.50169 | 0.51925 | 0.47401 | 0.50796 | 0.47919 | 0.51748 | 0.49157 | 0.50113 | 0.51858 | 1.23936 | 0.4869 | 0.48459 | 0.48992 | 0.5108 |
| ID17 | 0.47917 | 0.5234 | 0.48443 | 0.54176 | 0.50227 | 0.48702 | 0.49887 | 0.4886 | 0.50741 | 0.50517 | 0.46546 | 0.47839 | 0.49548 | 0.50171 | 0.49321 | 0.4869 | 1.22615 | 0.52644 | 0.50449 | 0.49429 |
| ID18 | 0.4836 | 0.50345 | 0.51095 | 0.50859 | 0.49319 | 0.51359 | 0.48804 | 0.4943 | 0.49715 | 0.50919 | 0.45598 | 0.50342 | 0.49489 | 0.47722 | 0.50227 | 0.48459 | 0.52644 | 1.24464 | 0.50961 | 0.49141 |
| ID19 | 0.48667 | 0.49719 | 0.50962 | 0.50899 | 0.49278 | 0.5061 | 0.5167 | 0.55387 | 0.49777 | 0.4955 | 0.48556 | 0.50723 | 0.52722 | 0.49665 | 0.49499 | 0.48992 | 0.50449 | 0.50961 | 1.24185 | 0.50224 |
| ID20 | 0.49606 | 0.50795 | 0.48744 | 0.49943 | 0.51179 | 0.50901 | 0.49606 | 0.49831 | 0.49095 | 0.49886 | 0.52301 | 0.49211 | 0.52301 | 0.49548 | 0.48647 | 0.5108 | 0.49429 | 0.49141 | 0.50224 | 1.25106 |

**Supplementary table 2.**
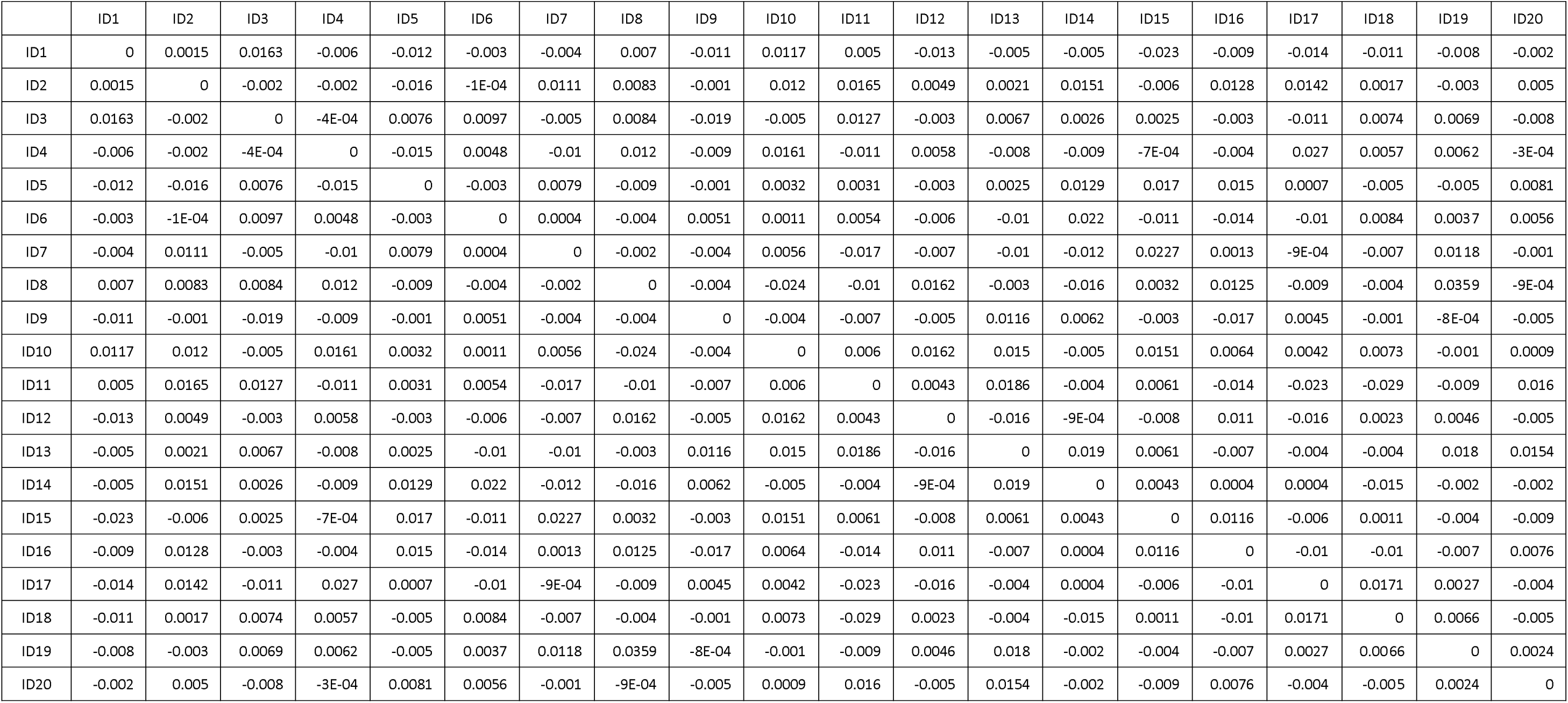
Kinship matrix calculated based on the method of Loiselle et al. (1995) using SPAGeDi. All elemental values on the diagonal are zero, and negative values are present in the matrix.

|  | ID1 | ID2 | ID3 | ID4 | ID5 | ID6 | ID7 | ID8 | ID9 | ID10 | ID11 | ID12 | ID13 | ID14 | ID15 | ID16 | ID17 | ID18 | ID19 | ID20 |
| --- | --- | --- | --- | --- | --- | --- | --- | --- | --- | --- | --- | --- | --- | --- | --- | --- | --- | --- | --- | --- |
| ID1 | 0 | 0.0015 | 0.0163 | -0.006 | -0.012 | -0.003 | -0.004 | 0.007 | -0.011 | 0.0117 | 0.005 | -0.013 | -0.005 | -0.005 | -0.023 | -0.009 | -0.014 | -0.011 | -0.008 | -0.002 |
| ID2 | 0.0015 | 0 | -0.002 | -0.002 | -0.016 | -1E-04 | 0.0111 | 0.0083 | -0.001 | 0.012 | 0.0165 | 0.0049 | 0.0021 | 0.0151 | -0.006 | 0.0128 | 0.0142 | 0.0017 | -0.003 | 0.005 |
| ID3 | 0.0163 | -0.002 | 0 | -4E-04 | 0.0076 | 0.0097 | -0.005 | 0.0084 | -0.019 | -0.005 | 0.0127 | -0.003 | 0.0067 | 0.0026 | 0.0025 | -0.003 | -0.011 | 0.0074 | 0.0069 | -0.008 |
| ID4 | -0.006 | -0.002 | -4E-04 | 0 | -0.015 | 0.0048 | -0.01 | 0.012 | -0.009 | 0.0161 | -0.011 | 0.0058 | -0.008 | -0.009 | -7E-04 | -0.004 | 0.027 | 0.0057 | 0.0062 | -3E-04 |
| ID5 | -0.012 | -0.016 | 0.0076 | -0.015 | 0 | -0.003 | 0.0079 | -0.009 | -0.001 | 0.0032 | 0.0031 | -0.003 | 0.0025 | 0.0129 | 0.017 | 0.015 | 0.0007 | -0.005 | -0.005 | 0.0081 |
| ID6 | -0.003 | -1E-04 | 0.0097 | 0.0048 | -0.003 | 0 | 0.0004 | -0.004 | 0.0051 | 0.0011 | 0.0054 | -0.006 | -0.01 | 0.022 | -0.011 | -0.014 | -0.01 | 0.0084 | 0.0037 | 0.0056 |
| ID7 | -0.004 | 0.0111 | -0.005 | -0.01 | 0.0079 | 0.0004 | 0 | -0.002 | -0.004 | 0.0056 | -0.017 | -0.007 | -0.01 | -0.012 | 0.0227 | 0.0013 | -9E-04 | -0.007 | 0.0118 | -0.001 |
| ID8 | 0.007 | 0.0083 | 0.0084 | 0.012 | -0.009 | -0.004 | -0.002 | 0 | -0.004 | -0.024 | -0.01 | 0.0162 | -0.003 | -0.016 | 0.0032 | 0.0125 | -0.009 | -0.004 | 0.0359 | -9E-04 |
| ID9 | -0.011 | -0.001 | -0.019 | -0.009 | -0.001 | 0.0051 | -0.004 | -0.004 | 0 | -0.004 | -0.007 | -0.005 | 0.0116 | 0.0062 | -0.003 | -0.017 | 0.0045 | -0.001 | -8E-04 | -0.005 |
| ID10 | 0.0117 | 0.012 | -0.005 | 0.0161 | 0.0032 | 0.0011 | 0.0056 | -0.024 | -0.004 | 0 | 0.006 | 0.0162 | 0.015 | -0.005 | 0.0151 | 0.0064 | 0.0042 | 0.0073 | -0.001 | 0.0009 |
| ID11 | 0.005 | 0.0165 | 0.0127 | -0.011 | 0.0031 | 0.0054 | -0.017 | -0.01 | -0.007 | 0.006 | 0 | 0.0043 | 0.0186 | -0.004 | 0.0061 | -0.014 | -0.023 | -0.029 | -0.009 | 0.016 |
| ID12 | -0.013 | 0.0049 | -0.003 | 0.0058 | -0.003 | -0.006 | -0.007 | 0.0162 | -0.005 | 0.0162 | 0.0043 | 0 | -0.016 | -9E-04 | -0.008 | 0.011 | -0.016 | 0.0023 | 0.0046 | -0.005 |
| ID13 | -0.005 | 0.0021 | 0.0067 | -0.008 | 0.0025 | -0.01 | -0.01 | -0.003 | 0.0116 | 0.015 | 0.0186 | -0.016 | 0 | 0.019 | 0.0061 | -0.007 | -0.004 | -0.004 | 0.018 | 0.0154 |
| ID14 | -0.005 | 0.0151 | 0.0026 | -0.009 | 0.0129 | 0.022 | -0.012 | -0.016 | 0.0062 | -0.005 | -0.004 | -9E-04 | 0.019 | 0 | 0.0043 | 0.0004 | 0.0004 | -0.015 | -0.002 | -0.002 |
| ID15 | -0.023 | -0.006 | 0.0025 | -7E-04 | 0.017 | -0.011 | 0.0227 | 0.0032 | -0.003 | 0.0151 | 0.0061 | -0.008 | 0.0061 | 0.0043 | 0 | 0.0116 | -0.006 | 0.0011 | -0.004 | -0.009 |
| ID16 | -0.009 | 0.0128 | -0.003 | -0.004 | 0.015 | -0.014 | 0.0013 | 0.0125 | -0.017 | 0.0064 | -0.014 | 0.011 | -0.007 | 0.0004 | 0.0116 | 0 | -0.01 | -0.01 | -0.007 | 0.0076 |
| ID17 | -0.014 | 0.0142 | -0.011 | 0.027 | 0.0007 | -0.01 | -9E-04 | -0.009 | 0.0045 | 0.0042 | -0.023 | -0.016 | -0.004 | 0.0004 | -0.006 | -0.01 | 0 | 0.0171 | 0.0027 | -0.004 |
| ID18 | -0.011 | 0.0017 | 0.0074 | 0.0057 | -0.005 | 0.0084 | -0.007 | -0.004 | -0.001 | 0.0073 | -0.029 | 0.0023 | -0.004 | -0.015 | 0.0011 | -0.01 | 0.0171 | 0 | 0.0066 | -0.005 |
| ID19 | -0.008 | -0.003 | 0.0069 | 0.0062 | -0.005 | 0.0037 | 0.0118 | 0.0359 | -8E-04 | -0.001 | -0.009 | 0.0046 | 0.018 | -0.002 | -0.004 | -0.007 | 0.0027 | 0.0066 | 0 | 0.0024 |
| ID20 | -0.002 | 0.005 | -0.008 | -3E-04 | 0.0081 | 0.0056 | -0.001 | -9E-04 | -0.005 | 0.0009 | 0.016 | -0.005 | 0.0154 | -0.002 | -0.009 | 0.0076 | -0.004 | -0.005 | 0.0024 | 0 |

**Supplementary table 3.** Normalized identical by state matrix calculated based on the method of Yang et al. (2011) using TASSEL. Negative values are present in the matrix.

|  | ID1 | ID2 | ID3 | ID4 | ID5 | ID6 | ID7 | ID8 | ID9 | ID10 | ID11 | ID12 | ID13 | ID14 | ID15 | ID16 | ID17 | ID18 | ID19 | ID20 |
| --- | --- | --- | --- | --- | --- | --- | --- | --- | --- | --- | --- | --- | --- | --- | --- | --- | --- | --- | --- | --- |
| ID1 | 0.9663 | -0.0114 | 0.0269 | -0.0070 | -0.0419 | 0.0120 | -0.0347 | -0.0132 | -0.0421 | 0.0419 | 0.0309 | -0.0231 | -0.0112 | 0.0234 | -0.0921 | -0.0454 | -0.0586 | -0.0057 | -0.0445 | 0.0281 |
| ID2 | -0.0114 | 0.9817 | 0.0173 | 0.0189 | -0.0432 | 0.0147 | -0.0341 | 0.0192 | 0.0623 | -0.0064 | 0.0124 | 0.0129 | 0.0051 | 0.0508 | 0.0303 | 0.0569 | 0.0410 | -0.0007 | -0.0283 | -0.0005 |
| ID3 | 0.0269 | 0.0173 | 1.0254 | -0.0152 | 0.0235 | 0.0302 | 0.0161 | 0.0116 | -0.0542 | -0.0025 | 0.0083 | -0.0278 | -0.0006 | -0.0435 | 0.0094 | 0.0182 | 0.0071 | -0.0312 | 0.0537 | 0.0266 |
| ID4 | -0.0070 | 0.0189 | -0.0152 | 0.9591 | -0.0053 | 0.0529 | -0.0231 | 0.0537 | -0.0637 | 0.0011 | -0.0551 | -0.0193 | -0.0010 | -0.0353 | -0.0146 | -0.0132 | 0.0746 | 0.0515 | -0.0035 | -0.0729 |
| ID5 | -0.0419 | -0.0432 | 0.0235 | -0.0053 | 0.9577 | -0.0163 | 0.0136 | -0.0236 | -0.0179 | -0.0227 | 0.0113 | 0.0318 | -0.0035 | 0.0482 | 0.0547 | 0.0006 | 0.0322 | -0.0172 | 0.0155 | 0.0015 |
| ID6 | 0.0120 | 0.0147 | 0.0302 | 0.0529 | -0.0163 | 1.1089 | -0.0059 | -0.0082 | -0.0076 | 0.0050 | 0.0025 | 0.0442 | -0.0336 | 0.0899 | -0.0469 | 0.0052 | -0.0605 | 0.0778 | -0.0341 | -0.0232 |
| ID7 | -0.0347 | -0.0341 | 0.0161 | -0.0231 | 0.0136 | -0.0059 | 0.9603 | -0.0002 | -0.0349 | 0.0044 | -0.0780 | -0.0052 | -0.0233 | -0.0803 | 0.0573 | -0.0118 | 0.0075 | -0.0182 | 0.0084 | 0.0228 |
| ID8 | -0.0132 | 0.0192 | 0.0116 | 0.0537 | -0.0236 | -0.0082 | -0.0002 | 1.0862 | -0.0351 | -0.0971 | -0.0429 | 0.0206 | 0.0198 | -0.0199 | 0.0046 | -0.0067 | 0.0413 | 0.0531 | 0.0861 | -0.0353 |
| ID9 | -0.0421 | 0.0623 | -0.0542 | -0.0637 | -0.0179 | -0.0076 | -0.0349 | -0.0351 | 0.9715 | -0.0193 | 0.0056 | -0.0300 | 0.0265 | 0.0400 | -0.0426 | -0.0005 | 0.0088 | -0.0316 | 0.0366 | -0.0015 |
| ID10 | 0.0419 | -0.0064 | -0.0025 | 0.0011 | -0.0227 | 0.0050 | 0.0044 | -0.0971 | -0.0193 | 1.0154 | 0.0083 | 0.0492 | -0.0002 | -0.0174 | 0.0652 | 0.0262 | -0.0553 | 0.0194 | 0.0201 | 0.0106 |
| ID11 | 0.0309 | 0.0124 | 0.0083 | -0.0551 | 0.0113 | 0.0025 | -0.0780 | -0.0429 | 0.0056 | 0.0083 | 1.0375 | 0.0162 | 0.0346 | 0.0184 | 0.0116 | 0.0192 | -0.0599 | -0.0262 | 0.0020 | 0.0407 |
| ID12 | -0.0231 | 0.0129 | -0.0278 | -0.0193 | 0.0318 | 0.0442 | -0.0052 | 0.0206 | -0.0300 | 0.0492 | 0.0162 | 1.0621 | -0.0709 | -0.0214 | 0.0104 | -0.0123 | 0.0115 | 0.0414 | -0.0026 | -0.0069 |
| ID13 | -0.0112 | 0.0051 | -0.0006 | -0.0010 | -0.0035 | -0.0336 | -0.0233 | 0.0198 | 0.0265 | -0.0002 | 0.0346 | -0.0709 | 1.0752 | 0.0614 | 0.0036 | 0.0598 | 0.0187 | -0.0058 | 0.0328 | 0.0190 |
| ID14 | 0.0234 | 0.0508 | -0.0435 | -0.0353 | 0.0482 | 0.0899 | -0.0803 | -0.0199 | 0.0400 | -0.0174 | 0.0184 | -0.0214 | 0.0614 | 1.0603 | -0.0232 | 0.0074 | -0.0148 | -0.0181 | 0.0462 | 0.0176 |
| ID15 | -0.0921 | 0.0303 | 0.0094 | -0.0146 | 0.0547 | -0.0469 | 0.0573 | 0.0046 | -0.0426 | 0.0652 | 0.0116 | 0.0104 | 0.0036 | -0.0232 | 0.9579 | 0.0330 | 0.0060 | 0.0246 | 0.0103 | -0.0204 |
| ID16 | -0.0454 | 0.0569 | 0.0182 | -0.0132 | 0.0006 | 0.0052 | -0.0118 | -0.0067 | -0.0005 | 0.0262 | 0.0192 | -0.0123 | 0.0598 | 0.0074 | 0.0330 | 0.9849 | -0.0292 | 0.0198 | -0.0151 | -0.0095 |
| ID17 | -0.0586 | 0.0410 | 0.0071 | 0.0746 | 0.0322 | -0.0605 | 0.0075 | 0.0413 | 0.0088 | -0.0553 | -0.0599 | 0.0115 | 0.0187 | -0.0148 | 0.0060 | -0.0292 | 0.9634 | 0.0142 | 0.0482 | -0.0253 |
| ID18 | -0.0057 | -0.0007 | -0.0312 | 0.0515 | -0.0172 | 0.0778 | -0.0182 | 0.0531 | -0.0316 | 0.0194 | -0.0262 | 0.0414 | -0.0058 | -0.0181 | 0.0246 | 0.0198 | 0.0142 | 0.9361 | -0.0014 | -0.0028 |
| ID19 | -0.0445 | -0.0283 | 0.0537 | -0.0035 | 0.0155 | -0.0341 | 0.0084 | 0.0861 | 0.0366 | 0.0201 | 0.0020 | -0.0026 | 0.0328 | 0.0462 | 0.0103 | -0.0151 | 0.0482 | -0.0014 | 0.9629 | 0.0200 |
| ID20 | 0.0281 | -0.0005 | 0.0266 | -0.0729 | 0.0015 | -0.0232 | 0.0228 | -0.0353 | -0.0015 | 0.0106 | 0.0407 | -0.0069 | 0.0190 | 0.0176 | -0.0204 | -0.0095 | -0.0253 | -0.0028 | 0.0200 | 1.0196 |

